# The gut bacterial community potentiates *Clostridioides difficile* infection severity

**DOI:** 10.1101/2022.01.31.478599

**Authors:** Nicholas A. Lesniak, Alyxandria M. Schubert, Kaitlyn J. Flynn, Jhansi L. Leslie, Hamide Sinani, Ingrid L. Bergin, Vincent B. Young, Patrick D. Schloss

**Affiliations:** Department of Microbiology and Immunology, University of Michigan, Ann Arbor, MI; Division of Infectious Diseases, Department of Internal Medicine, University of Michigan Medical School, Ann Arbor, MI; Unit for Laboratory Animal Medicine, University of Michigan, Ann Arbor, MI; Department of Medicine, Division of International Health and Infectious Diseases, University of Virginia School of Medicine, Charlottesville, Virginia, USA

## Abstract

The severity of *Clostridioides difficile* infections (CDI) has increased over the last few decades. Patient age, white blood cell count, creatinine levels as well as *C. difficile* ribotype and toxin genes have been associated with disease severity. However, it is unclear whether there is an association between members of the gut microbiota and disease severity. The gut microbiota is known to interact with *C. difficile* during infection. Perturbations to the gut microbiota are necessary for *C. difficile* to colonize the gut. The gut microbiota can inhibit *C. difficile* colonization through bile acid metabolism, nutrient consumption and bacteriocin production. Here we sought to demonstrate that members of the gut bacterial communities can also contribute to disease severity. We derived diverse gut communities by colonizing germ-free mice with different human fecal communities. The mice were then infected with a single *C. difficile* ribotype 027 clinical isolate which resulted in moribundity and histopathologic differences. The variation in severity was associated with the human fecal community that the mice received. Generally, bacterial populations with pathogenic potential, such as *Escherichia*, *Helicobacter*, and *Klebsiella*, were associated with more severe outcomes. Bacterial groups associated with fiber degradation, bile acid metabolism and lantibiotic production, such as *Anaerostipes* and *Coprobacillus*, were associated with less severe outcomes. These data indicate that, in addition to the host and *C. difficile*, populations of gut bacteria can influence CDI disease severity.

**Importance:** *Clostridioides difficile* colonization can be asymptomatic or develop into an infection, ranging in severity from mild diarrhea to toxic megacolon, sepsis, and death. Models that predict severity and guide treatment decisions are based on clinical factors and *C. difficile* characteristics. Although the gut microbiome plays a role in protecting against CDI, its effect on CDI disease severity is unclear and has not been incorporated into disease severity models. We demonstrated that variation in the microbiome of mice colonized with human feces yielded a range of disease outcomes. These results revealed groups of bacteria associated with both severe and mild *C. difficile* infection outcomes. Gut bacterial community data from patients with CDI could improve our ability to identify patients at risk of developing more severe disease and improve interventions which target *C. difficile* and the gut bacteria to reduce host damage.

## Introduction

*Clostridioides difficile* infections (CDI) have increased in incidence and severity since *C. difficile* was first identified as the cause of antibiotic-associated pseudomembranous colitis (1). CDI disease severity can range from mild diarrhea to toxic megacolon and death. The Infectious Diseases Society of America (IDSA) and Society for Healthcare Epidemiology of America (SHEA) guidelines define severe CDI in terms of a white blood cell count greater than 15,000 cells/mm^3^ and/or a serum creatinine greater than 1.5 mg/dL. Patients who develop shock or hypotension, ileus, or toxic megacolon are considered to have fulminant CDI (2). Since these measures are CDI outcomes, they have limited ability to predict risk of severe CDI when the infection is first detected. Schemes have been developed to score a patient’s risk for severe CDI outcomes based on clinical factors but have not been robust for broad application (3). Thus, we have limited ability to prevent patients from developing severe CDI.

Missing from CDI severity prediction models are the effects of the indigenous gut bacteria. *C. difficile* interacts with the gut community in many ways. The indigenous bacteria of a healthy intestinal community provide a protective barrier preventing *C. difficile* from infecting the gut. A range of mechanisms can disrupt this barrier, including antibiotics, medications, or dietary changes, and lead to increased susceptibility to CDI (4–6). Once *C. difficile* overcomes the protective barrier and colonizes the intestine, the indigenous bacteria can either promote or inhibit *C. difficile* through producing molecules or modifying the environment (7, 8). Bile acids metabolized by the gut bacteria can inhibit *C. difficile* growth and affect toxin production (9, 10). Bacteria in the gut also can compete more directly with *C. difficile* through antibiotic production or nutrient consumption (11–13). While the relationship between the gut bacteria and *C. difficile* has been established, the effect the gut bacteria can have on CDI disease severity is unclear.

Recent studies have demonstrated that when mice with diverse microbial communities were challenged with a high-toxigenic strain resulted in varied disease severity (14) and when challenged with a low-toxigenic strain members of the gut microbial community associated with variation in colonization (15). Here, we sought to further elucidate the relationship between members of the gut bacterial community and CDI disease severity when challenged with a high-toxigenic strain, *C. difficile* ribotype 027 (RT027). We hypothesized that since specific groups of gut bacteria affect the metabolism of *C. difficile* and its infection dynamics, we can also identify groups of bacteria that affect the disease severity of the infection. To test this hypothesis, we colonized germ-free C57BL/6 mice with human fecal samples to create varied gut communities. We then challenged the mice with *C. difficile* RT027 and followed the mice for the development of severe outcomes of moribundity and histopathologic cecal tissue damage. Since the murine host and *C. difficile* isolate were the same and only the gut community varied, the variation in disease severity we observed was attributable to the gut microbiome.

## Results

### *C. difficile* is able to infect germ-free mice colonized with human fecal microbial communities without antibiotics

To produce gut microbiomes with greater variation than those found in conventional mouse colonies, we colonized germ-free mice with bacteria from human feces (16). We inoculated germ-free C57BL/6 mice with homogenized feces from each of 15 human fecal samples via oral gavage. These human fecal samples were selected because they represented diverse community structures based on community clustering (17). The gut communities were allowed to equilibrate for two weeks post-inoculation (18). We then surveyed the bacterial members of the gut communities by 16S rRNA gene sequencing of murine fecal pellets (Figure 1A). The bacterial communities from each mouse grouped more closely to those communities from mice that received the same human fecal donor community than to the mice who received a different human fecal donor community (Figure 1B). The communities were primarily composed of populations of *Clostridia*, *Bacteroidia*, *Erysipelotrichia*, *Bacilli*, and *Gammaproteobacteria*. However, the gut bacterial communities of each donor group of mice harbored unique relative abundance distributions of the shared bacterial classes.

**Figure 1.**
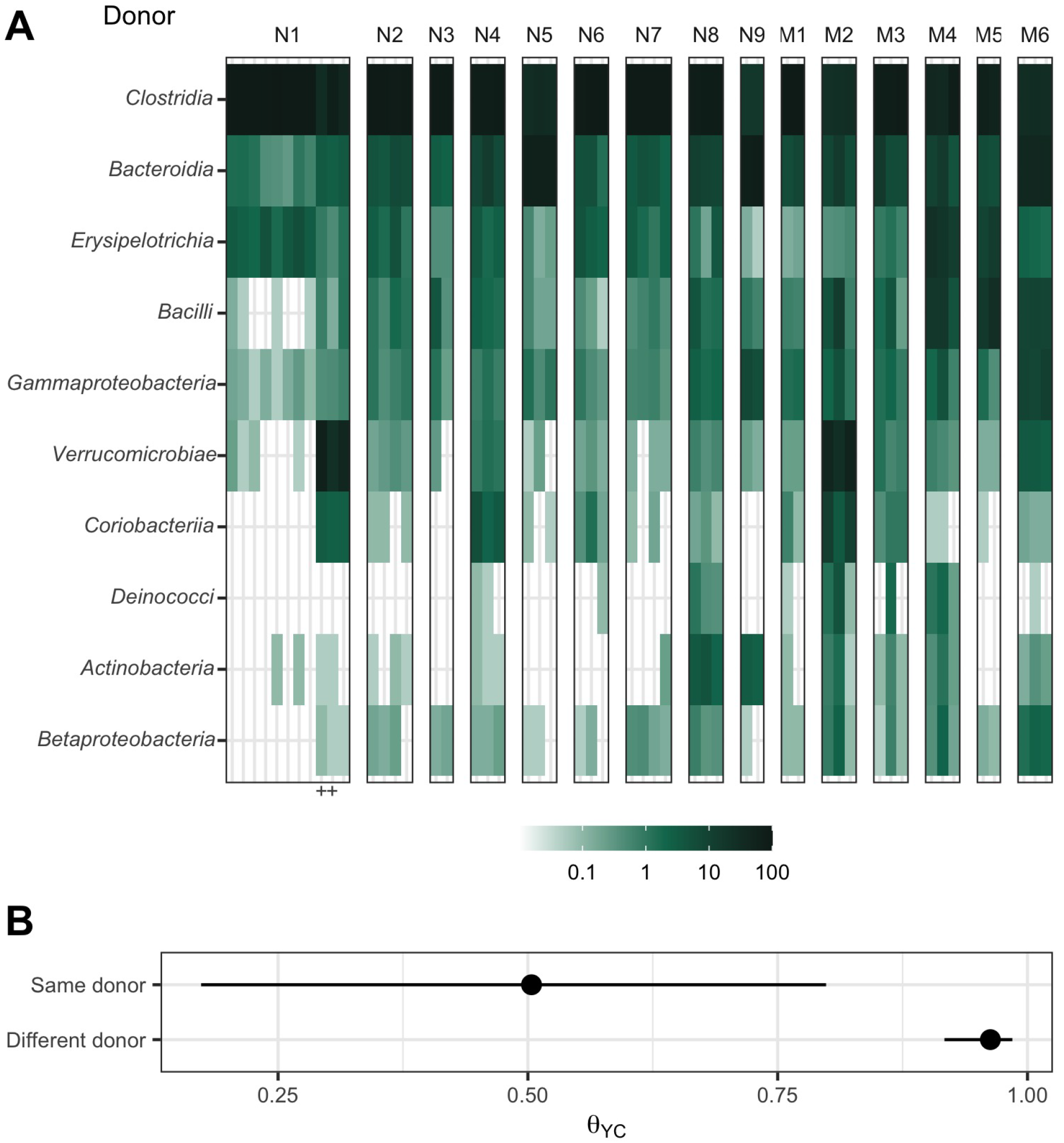
Human fecal microbial communities established diverse gut bacterial communities in germ-free mice. (A) Relative abundances of the 10 most abundant bacterial classes observed in the feces of previously germ-free C57Bl/6 mice 14 days post-colonization with human fecal samples (i.e., day 0 relative to *C. difficile* challenge). Each column of abundances represents an individual mouse. Mice that received the same donor feces are grouped together and labeled above with a letter (N for non-moribund mice and M for moribund mice) and number (ordered by mean histopathologic score of the donor group). + indicates the mice which did not have detectable *C. difficile* CFU (Figure 2). (B) Median (points) and interquartile range (lines) of *β*-diversity (*θ*_YC_) between an individual mouse and either all others which were inoculated with feces from the same donor or from a different donor. The *β*-diversity among the same donor comparison group was significantly less than the *β*-diversity of the different donor group (*P* < 0.05, calculated by Wilcoxon rank sum test).

Next, we tested this set of mice with their human-derived gut microbial communities for susceptibility to *C. difficile* infection. A typical mouse model of CDI requires pre-treatment of conventional mice with antibiotics, such as clindamycin, to become susceptible to *C. difficile* colonization (19, 20). However, we wanted to avoid modifying the gut communities with an antibiotic to maintain their unique microbial compositions and ecological relationships. Since some of these communities came from people at increased risk of CDI, such as recent hospitalization or antibiotic use (17), we tested whether *C. difficile* was able to infect these mice without an antibiotic perturbation. We hypothesized that *C. difficile* would be able to colonize the mice who received their gut communities from a donor with a perturbed community. Mice were challenged with 10^3^ *C. difficile* RT027 clinical isolate spores. The mice were followed for 10 days post-challenge, and their stool was collected and plated for *C. difficile* colony forming units (CFU) to determine the extent of the infection. Surprisingly, communities from all donors were able to be colonized (Figure 2). Two mice were able to resist *C. difficile* colonization, both received their community donor N1, which may be attributed to experimental variation since this group also had more mice. By colonizing germ-free mice with different human fecal communities, we were able to generate diverse gut communities in mice, which were susceptible to *C. difficile* infection without further modification of the gut community.

**Figure 2.**
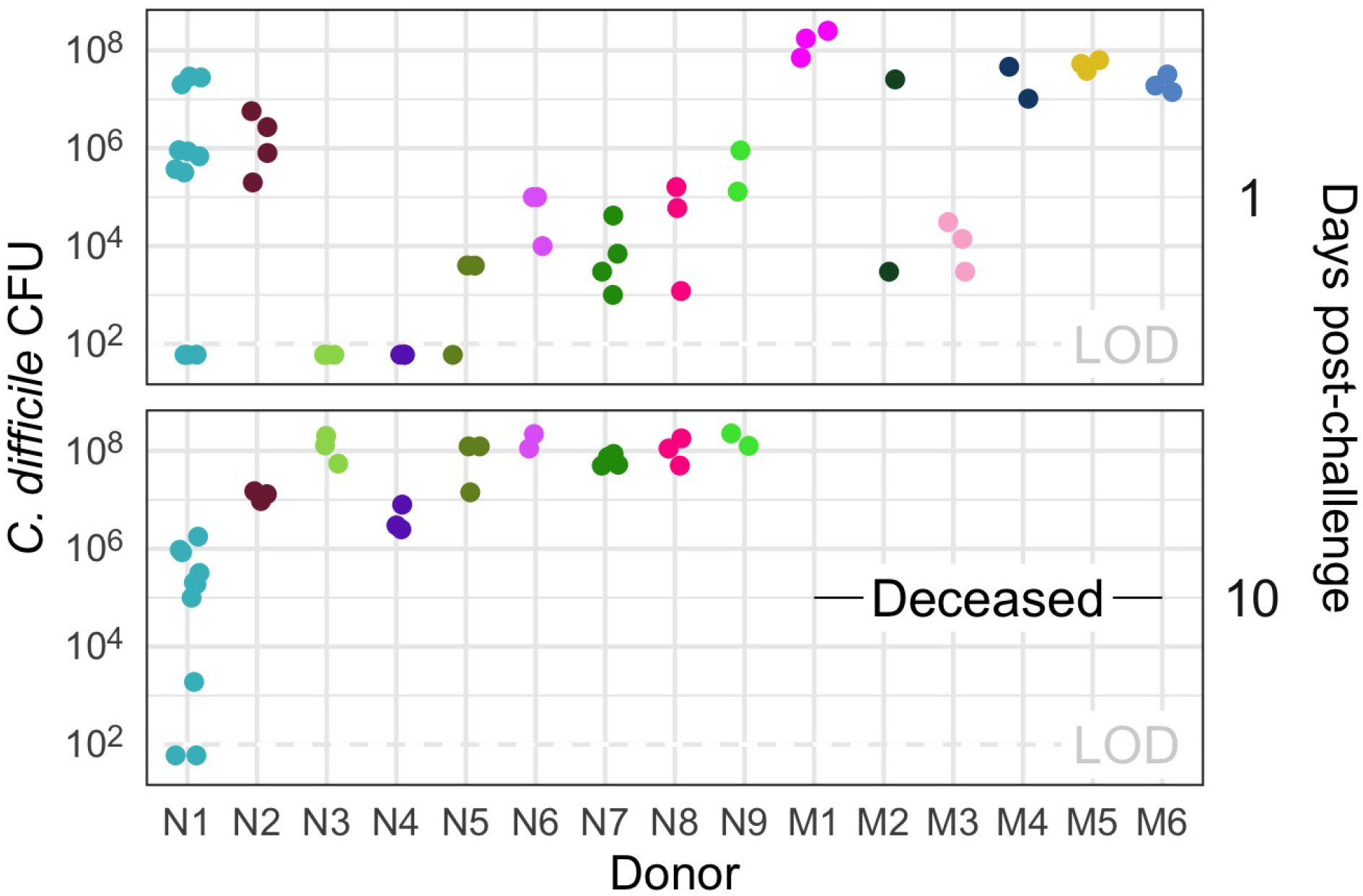
All donor groups resulted in *C. difficile* infection but with different outcomes. *C. difficile* CFU per gram of stool was measured the day after challenge with 10^3^ *C. difficile* RT027 clinical isolate 431 spores and at the end of the experiment, 10 days post-challenge. Each point represents an individual mouse. Mice are grouped by donor and labeled by the donor letter (N for non-moribund mice and M for moribund mice) and number (ordered by mean histopathologic score of the donor group). Points are colored by donor group. Mice from donor groups N1 through N6 succumbed to the infection prior to day 10 and were not plated on day 10 post-challenge. LOD = Limit of detection.

### Infection severity varies by initial community

After we challenged the mice with *C. difficile*, we investigated the outcome from the infection and its relationship to the initial community. We followed the mice for 10 days post-challenge for colonization density, toxin production, and mortality. Seven mice, from Donors N1, N3, N4, and N5, were not colonized at detectable levels on the day after *C. difficile* challenge but were infected (>10^6^) by the end of the experiment. All mice that received their community from Donor M1 through M6 succumbed to the infection and became moribund within 3 days post-challenge. The remaining mice, except the uninfected Donor N1 mice, maintained *C. difficile* infection through the end of the experiment (Figure 2). At 10 days post-challenge, or earlier for the moribund mice, mice were euthanised and fecal material were assayed for toxin activity and cecal tissue was collected and scored for histopathologic signs of disease (Figure 3). Overall, there was greater toxin activity detected in the stool of the moribund mice (*P* = 0.003). However, when looking at each group of mice, we observed a range in toxin activity for both the moribund and non-moribund mice (Figure 3A). Non-moribund mice from Donors N2 and N5 through N9 had comparable toxin activity as the moribund mice. Additionally, not all moribund mice had toxin activity detected in their stool. Next, we examined the cecal tissue for histopathologic damage. Moribund mice had high levels of epithelial damage, tissue edema, and inflammation (Figure S1) similar to previously reported histopathologic findings for *C. difficile* RT027 (21). As observed with toxin activity, the moribund mice had higher histopathologic scores than the non-moribund mice (*P* < 0.001). However, unlike the toxin activity, all moribund mice had consistently high histopathologic summary scores (Figure 3B). The non-moribund mice, Donor groups N1 through N9, had a range in tissue damage from none detected to similar levels as the moribund mice, which grouped by community donor. Together, the toxin activity, histopathologic score, and moribundity showed variation across the donor groups but were largely consistent within each donor group.

**Figure 3.**
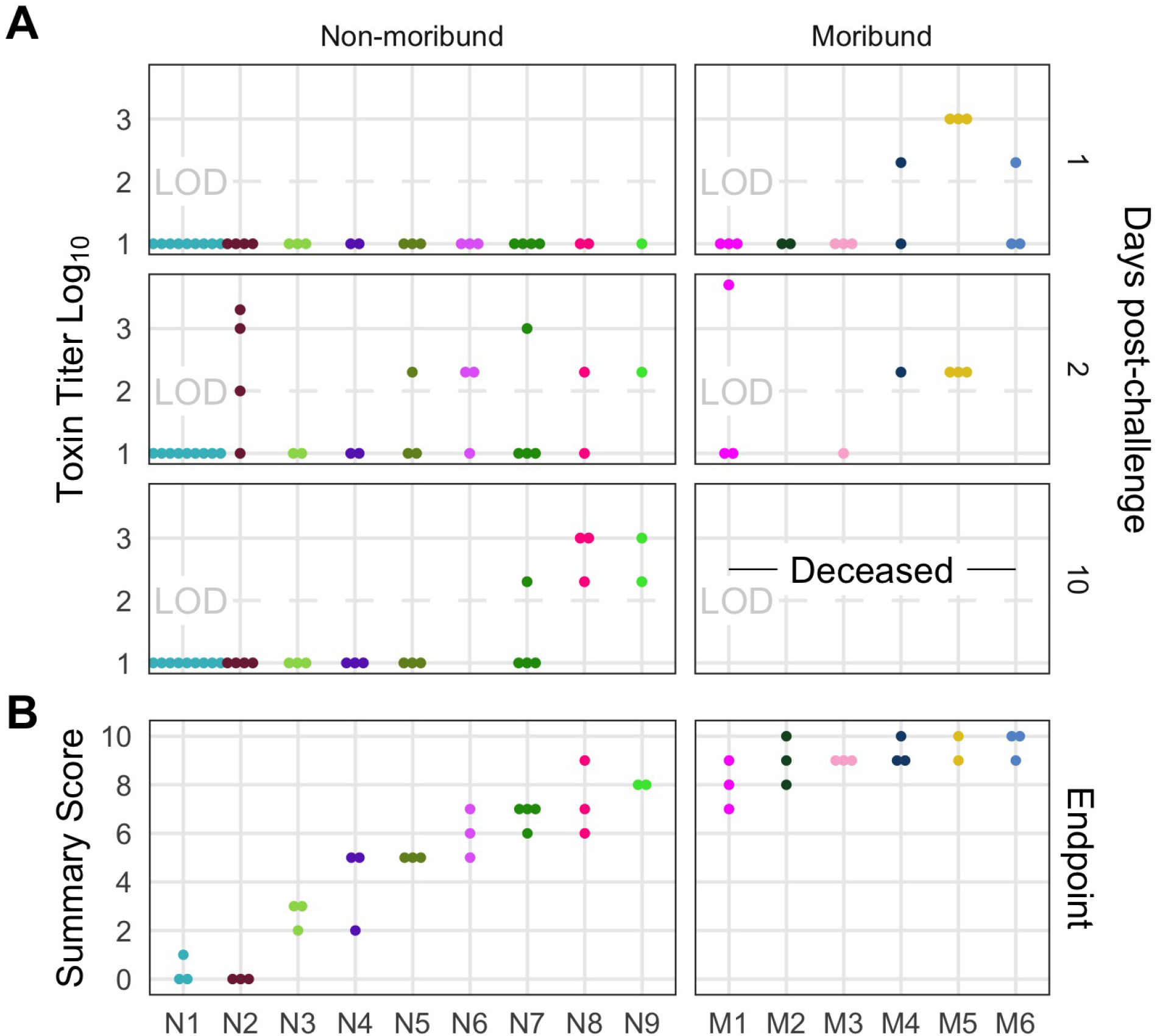
Histopathologic score and toxin activity varied across donor groups. (A) Fecal toxin activity was detected in some mice post *C. difficile* challenge in both moribund and non-moribund mice. (B) Cecum scored for histopathologic damage from mice at the end of the experiment. Samples were collected for histopathologic scoring on day 10 post-challenge for non-moribund mice or the day the mouse succumbed to the infection for the moribund group (day 2 or 3 post-challenge). Each point represents an individual mouse. Mice are grouped by donor and labeled by the donor letter (N for non-moribund mice and M for moribund mice) and number (ordered by mean histopathologic score of the donor group). Points are colored by donor group. Mice in group N1 that have a summary score of 0 are the mice which did not have detectable *C. difficile* CFU (Figure 2). Missing points are from mice that had insufficient fecal sample collected for assaying toxin or cecum for histopathologic scoring. LOD = Limit of detection.

### Microbial community members explain variation in CDI severity

We next interrogated the bacterial communities at the time of *C. difficile* challenge (day 0) for their relationship to infection outcomes using linear discriminant analysis (LDA) effect size (LEfSe) analysis to identify individual bacterial populations that could explain the variation in disease severity. We split the mice into groups by severity level based on their moribundity and histopathologic score. We dichotomized the histopathologic scores into high and low groups by splitting on the median score of 5. This analysis revealed 20 genera that were significantly different by the disease severity (Figure 4A). Bacterial genera *Turicibacter*, *Streptococcus*, *Staphylococcus*, *Pseudomonas*, *Phocaeicola*, *Parabacteroides*, *Bacteroides*, and *Escherichia/Shigella* were detected at higher relative abundances in the mice that became moribund. Populations of *Anaerotignum*, *Coprobacillus*, *Enterocloster*, and *Murimonas* were more abundant in the non-moribund mice that would develop only low intestinal injury. To understand the role of toxin activity in disease severity, we applied LEfSe to identify the genera most likely to explain the differences between the presence and absence of detected toxin activity (Figure 4B). Many genera that associated with the presence of toxin were also associated with moribundity, such as populations of *Escherichia/Shigella* and *Bacteroides*. Likewise, there were genera such as *Anaerotignum*, *Enterocloster*, and *Murimonas* that were associated with no detected toxin that also exhibited greater relative abundance in communities from non-moribund mice with a low histopathologic score. Lastly, we tested for correlations between the endpoint relative abundances of bacterial operational taxonomic units (OTUs) and the histopathologic summary score (Figure 4C). The endpoint relative abundance of *Bacteroides* was positively correlated with histopathologic score, as its day 0 relative abundance did with disease severity (Figure 4A). Populations of *Klebsiella* and *Prevotellaceae* were positively correlated with the histopathologic score and were increased in the group of mice with detectable toxin. This analysis identified bacterial genera that were associated with the variation in moribundity, histopathologic score, and toxin.

**Figure 4.**
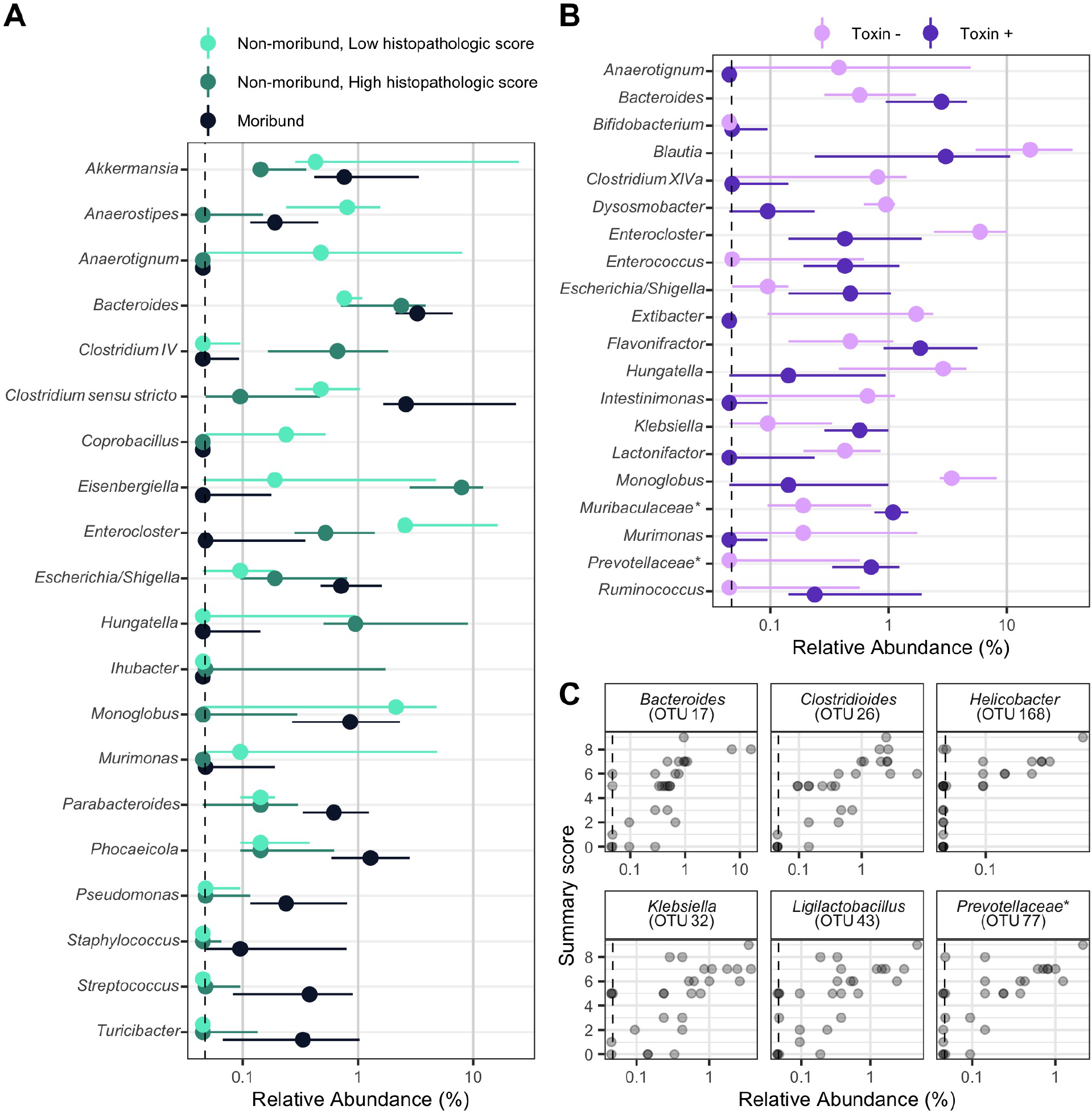
Individual fecal bacterial community members of the murine gut associated with *C .difficile* infection outcomes. (A and B) Relative abundance of genera at the time of *C. difficile* challenge (Day 0) that varied significantly by the moribundity and histopathologic summary score or detected toxin by LEfSe analysis. Median (points) and interquartile range (lines) are plotted. Genera are ordered alphabetically to ease comparisons across analyses. (A) Relative abundances were compared across infection outcome of moribund (colored black) or non-moribund with either a high histopathologic score (score greater than the median score of 5, colored green) or a low histopathologic summary score (score less than the median score of 5, colored light green). (B) Relative abundances were compared between mice which toxin activity was detected (Toxin +, colored dark purple) and which no toxin activity was detected (Toxin -, colored light purple). (C) Endpoint bacterial OTUs correlated with histopathologic summary score. Each individual mouse is plotted (transparent gray point). Spearman’s correlations were statistically significant after Benjamini-Hochberg correction for multiple comparisons. All bacterial groups are ordered alphabetically. * indicates that the bacterial group was unclassified at lower taxonomic classification ranks.

We next determined whether, collectively, bacterial community membership and relative abundance could be predictive of the CDI disease outcome. We trained random forest models with bacterial community relative abundance data from the day of colonization at each taxonomic rank to predict toxin, moribundity, and day 10 post-challenge histopathologic summary score. For predicting if detectable toxin would be produced, microbial populations aggregated by phylum rank classification performed similarly as models using lower taxonomic ranks (AUROC = 0.83, Figure S2). *C. difficile* was more likely to produce detectable toxin when the community infected had less abundant populations of *Verrucomicrobia* and *Campilobacterota* and had more abundant populations of *Proteobacteria* (Figure 5A). Next, we assessed the ability of the community to predict moribundity. Bacteria grouped by class rank classification was sufficient to predict which mice would succumb to the infection before the end of the experiment (AUROC = 0.91, Figure S2). The features with the greatest effect showed that communities with greater populations of bacteria belonging to *Bacilli* and *Firmicutes* and reduced populations of *Erysipelotrichia* were more likely to result in moribundity (Figure 5B). Only one other class of bacteria was decreased in moribund mice, a group of unclassified *Clostridia*. Lastly, the relative abundances of genera were able to predict a high or low histopathologic score (histopathologic scores were dichotomized as in previous analysis, AUROC = 0.99, Figure S2). No genera had a significantly greater effect on the model performance than any others, indicating the model was reliant on many genera for the correct prediction. The model used some of the genera identified in the LEfSe analysis, such as *Coprobacillus*, *Anaerostipes*, and *Hungatella*. Communities with greater abundances of *Hungatella*, *Eggerthella*, *Bifidobacterium*, *Duncaniella* and *Neisseria* were more likely to have high histopathologic scores. These models have shown that the relative abundance of bacterial populations and their relationship to each other could be used to predict the variation in moribundity, histopathologic score, and detectable toxin of CDI.

**Figure 5.**
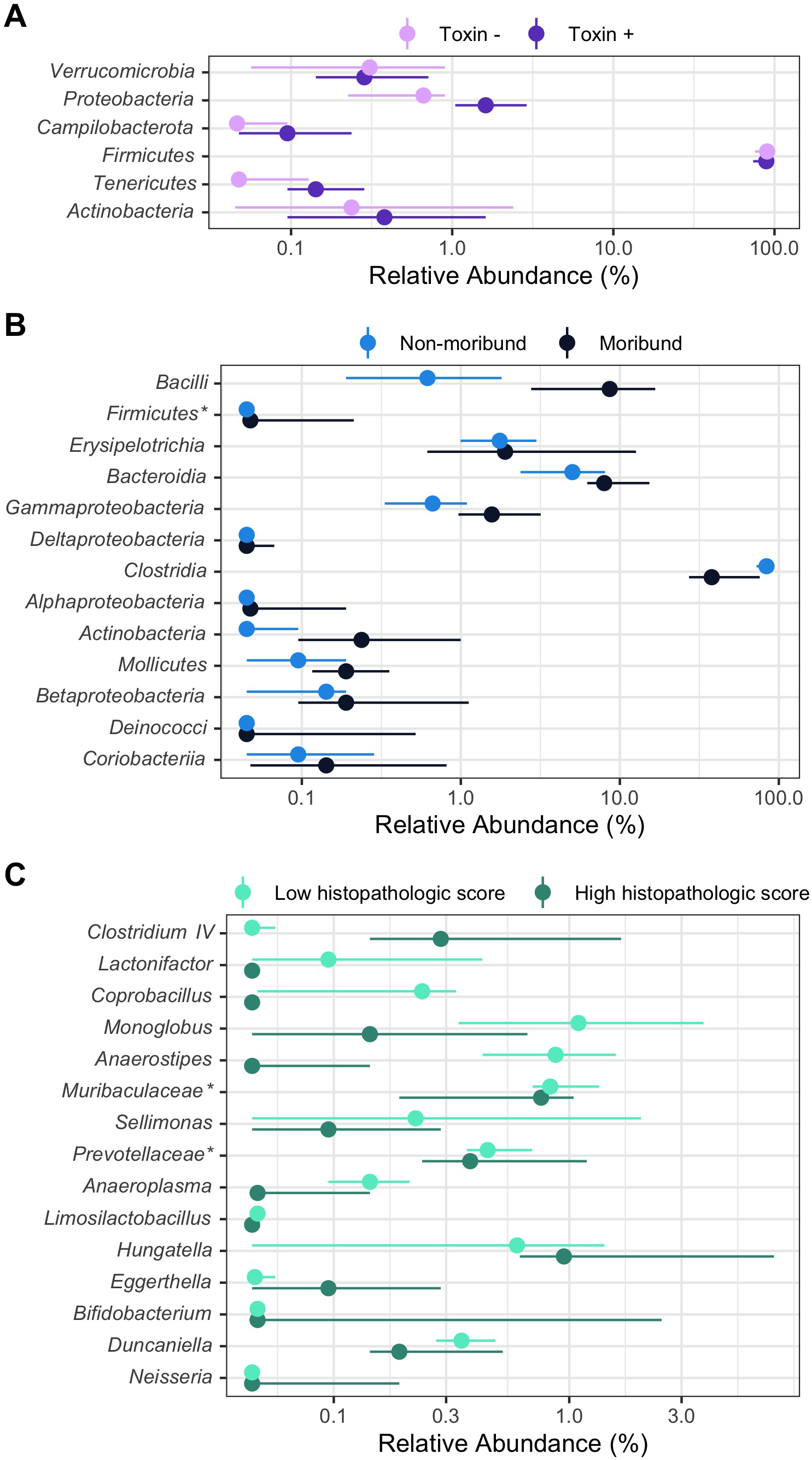
Fecal bacterial community members of the murine gut at the time of *C. difficile* infection predicted outcomes of the infection. On the day of infection (Day 0), bacterial community members grouped by different classification rank were modeled with random forest to predict the infection outcome. The models used the highest taxonomic classification rank that performed as well as the lower ranks. Median (solid points) and interquartile range (lines) of the group relative abundance are plotted. Bacterial groups are ordered by their importance to the model; taxonomic group at the top of the plot had the greatest decrease in performance when its relative abundances were permuted. * indicates that the bacterial group was unclassified at lower taxonomic classification ranks. (A) Bacterial members grouped by phyla predicted which mice would have toxin activity detected at any point throughout the infection (Toxin +, dark purple). (B) Bacterial members grouped by class predicted which mice would become moribund (dark blue). (C) Bacterial members grouped by genera predicted if the mice would have a high (score greater than the median score of 5, colored dark green) or low (score less than the median score of 5, colored light green) histopathologic summary score.

## Discussion

Challenging mice colonized with different human fecal communities with *C. difficile* RT027 demonstrated that variation in members of the gut microbiome affects *C. difficile* infection disease severity. Our analysis revealed an association between the relative abundance of bacterial community members and disease severity. Previous studies investigating the severity of CDI disease involving the microbiome have had limited ability to interrogate this relationship between the microbiome and disease severity. Studies that have used clinical data have limited ability to control variation in the host, microbiome or *C. difficile* ribotype (22). Murine experiments typically use a single mouse colony and different *C. difficile* ribotypes to create severity differences (23). Recently, our group has begun uncovering the effect microbiome variation has on *C. difficile* infection. We showed the variation in the bacterial communities between mice from different mouse colonies resulted in different clearance rates of *C. difficile* (15). We also showed varied ability of mice to spontaneously eliminate *C. difficile* infection when they were treated with different antibiotics prior to *C. difficile* challenge (24). Overall, the results presented here have demonstrated that the gut bacterial community contributed to the severity of *C. difficile* infection.

*C. difficile* can lead to asymptomatic colonization or infections with severity ranging from mild diarrhea to death. Physicians use classification tools to identify patients most at risk of developing a severe infection using white blood cell counts, serum albumin level, or serum creatinine level (2, 25, 26). Those levels are driven by the activities in the intestine (27). Research into the drivers of this variation have revealed factors that make *C. difficile* more virulent. Strains are categorized for their virulence by the presence and production of the toxins TcdA, TcdB, and binary toxin and the prevalence in outbreaks, such as ribotypes 027 and 078 (19, 28–31). However, other studies have shown that disease is not necessarily linked with toxin production (32) or the strain (33). Furthermore, there is variation in the genome, growth rate, sporulation, germination, and toxin production in different isolates of a strain (34–37). This variation may help explain why severe CDI prediction tools often miss identifying many patients with CDI that will develop severe disease (3, 23, 38, 39). Therefore, it is necessary to gain a full understanding of all factors contributing to disease variation to improve our ability to predict severity.

The state of the gut bacterial community determines the ability of *C. difficile* to colonize and persist in the intestine. *C. difficile* is unable to colonize an unperturbed healthy murine gut community and is only able to become established after a perturbation (20). Once colonized, the different communities lead to different metabolic responses and dynamics of the *C. difficile* population (8, 24, 40). Gut bacteria metabolize primary bile acids into secondary bile acids (41, 42). The concentration of these bile acids affects germination, growth, toxin production and biofilm formation (9, 10, 43, 44). Members of the bacterial community also affect other metabolites *C. difficile* utilizes. *Bacteroides thetaiotaomicron* produce sialidases which release sialic acid from the mucosa for *C. difficile* to utilize (45, 46). The nutrient environment affects toxin production (47). Thus, many of the actions of the gut bacteria modulate *C. difficile* in ways that could affect the infection and resultant disease.

A myriad of studies have explored the relationship between the microbiome and CDI disease. Studies examining difference in disease often use different *C. difficile* strains or ribotypes in mice with similar microbiota as a proxy for variation in disease, such as strain 630 for non-severe and RT027 for severe (19, 28, 29, 48). Studies have also demonstrated variation in infection through tapering antibiotic dosage (20, 24, 49) or by reducing the amount of *C. difficile* cells or spores used for the challenge (19, 49). These studies often either lack variation in the initial microbiome or have variation in the *C. difficile* infection itself, confounding any association between variation in severity and the microbiome. Recent studies have shown variation in the initial microbiome, via different murine colonies or colonizing germ-free mice with human feces, that were challenged with *C. difficile* resulted in varied outcomes of the infection (14, 15).

Our data have demonstrated gut bacterial relative abundances associate with variation in toxin production, histopathologic scoring of the cecal tissue and mortality. This analysis revealed populations of *Akkermansia*, *Anaerostipes*, *Coprobacillus*, *Enterocloster*, *Lactonifactor*, and *Monoglobus* were more abundant in the microbiome of non-moribund mice which had low histopathologic scores and no detected toxin. The protective role of these genera are supported by previous studies. *Coprobacillus*, *Lactonifactor*, and *Monoglobus* have been shown to be involved in dietary fiber fermentation and associated with healthy communities (50–53). *Anaerostipes* and *Coprobacillus*, which produce short chain fatty acids, have been associated with healthy communities (54–56). Furthermore, *Coprobacillus*, which was abundant in mice with low histopathologic scores but rare in all other mice, has been shown to contain a putative type I lantibiotic gene cluster and inhibit *C. difficile* colonization (57–59). *Akkermansia* and *Enterocloster* were also identified as more abundant in mice which had a low histopathologic scores but have contradictory supporting evidence in the current literature. In our data, *Akkermansia* was most abundant in the non-moribund mice with low histopathologic scores but there were some moribund mice which had increased populations of *Akkermansia*. This could be attributed to either a more protective mucus layer was present inhibiting colonization (59, 60) or mucus consumption by *Akkermansia* could have been crossfeeding *C. difficile* or exposing a niche for *C. difficile* (61–63). Similarly, *Enterocloster* was more abundant and associated with low histopathologic scores. It has been associated with healthy populations and has been used to mono-colonize germ-free mice to reduce the ability of *C. difficile* to colonize (64, 65). However, *Enterocloster* has also been involved in infections, such as bacteremia (66, 67). These data have exemplified populations of bacteria that have the potential to be either protective or harmful. Thus, the disease outcome is not likely based on the abundance of individual populations of bacteria, rather it is the result of the interactions of the community.

The groups of bacteria that were associated with either a higher histopathologic score or moribundity are members of the indigenous gut community that also have been associated with disease, often referred to as opportunistic pathogens. Many of the populations with pathogenic potential that associated with worse outcomes are also facultative anaerobes. *Enterococcus*, *Klebsiella*, *Shigella/Escherichia*, *Staphylococcus*, and *Streptococcus* have been shown to expand after antibiotic use (17, 68, 69) and are commonly detected in CDI cases (70–73). In addition to these populations, *Eggerthella*, *Prevotellaceae* and *Helicobacter*, which associated with worse outcomes, have also been associated with intestinal inflammation (74–76). Recently, *Helicobacter hepaticus* was shown to be sufficient to cause susceptibility to CDI in IL-10 deficient C57BL/6 mice (77). In our experiments, when *Helicobacter* was present, the infection resulted in a high histopathologic score (Figure 4C). While we did not use IL-10 deficient mice, it is possible the bacterial community or host response are similarly modified by *Helicobacter*, allowing *C. difficile* infection and host damage. Aside from *Helicobacter*, these groups of bacteria that associated with more severe outcomes did not have a conserved association between their relative abundance and the disease severity across all mice.

Since we observed groups of bacteria that were associated with less severe disease it may be appropriate to apply the damage-response framework for microbial pathogenesis to CDI (78, 79). This framework posits that disease is not driven by a single entity, rather it is an emergent property of the responses of the host immune system, infecting microbe, *C. difficile*, and the indigenous microbes at the site of infection. In the first set of experiments, we used the same host background, C57BL/6 mice, the same infecting microbe, *C. difficile* RT027 clinical isolate 431, with different gut bacterial communities. The bacterial groups in those communities were often present in both moribund and non-moribund and across the range of histopathologic scores. Thus, it was not merely the presence of the bacteria but their activity in response to the other microbes and host which affect the extent of the host damage. Additionally, while each mouse and *C. difficile* population had the same genetic background, they too were reacting to the specific microbial community. Disease severity is driven by the cumulative effect of the host immune response and the activity of *C. difficile* and the gut bacteria. *C. difficile* drives host damage through the production of toxin. The gut microbiota can modulate host damage through the balance of metabolic and competitive interactions with *C. difficile*, such as bacteriocin production or mucin degradation, and interactions with the host, such as host mucus glycosylation or intestinal IL-33 expression (14, 80). For example, low levels of mucin degradation can provide nutrients to other community members producing a diverse non-damaging community (81). However, if mucin degradation becomes too great it reduces the protective function of the mucin layer and exposes the epithelial cells. This over-harvesting can contribute to the host damage due to other members producing toxin. Thus, the resultant intestinal damage is the balance of all activities in the gut environment. Host damage is the emergent property of numerous damage-response curves, such as one for host immune response, one for *C. difficile* activity and another for microbiome community activity, each of which are a composite curve of the individual activities from each group, such as antibody production, neutrophil infiltration, toxin production, sporulation, fiber and mucin degradation. Therefore, while we have identified populations of interest, it may be necessary to target multiple types of bacteria to reduce the community interactions contributing to host damage.

Here we have shown several bacterial groups and their relative abundances associated with variation in CDI disease severity. Further understanding how the microbiome affects severity in patients could reduce the amount of adverse CDI outcomes. When a patient is diagnosed with CDI, the gut community composition, in addition to the traditionally obtained clinical information, may improve our severity prediction and guide prophylactic treatment. Treating the microbiome at the time of diagnosis, in addition to *C. difficile*, may prevent the infection from becoming more severe.

## Materials and Methods

### Animal care

6- to 13-week old male and female germ-free C57BL/6 were obtained from a single breeding colony in the University of Michigan Germ-free Mouse Core. Mice (N1 n=11, N2 n=7, N3 n=3, N4 n=3, N5 n=3, N6 n=3, N7 n=7, N8 n=3, N9 n=2, M1 n=3, M2 n=3, M3 n=3, M4 n=3, M5 n=7, M6 n=3) were housed in cages of 2-4 mice per cage and maintained in germ-free isolators at the University of Michigan germ-free facility. All mouse experiments were approved by the University Committee on Use and Care of Animals at the University of Michigan.

### *C. difficile* experiments

Human fecal samples were obtained as part of Schubert *et al*. and selected based on community clusters (17) to result in diverse community structures. Feces were homogenized by mixing 200 mg of sample with 5 ml of PBS. Mice were inoculated with 100 *μ*l of the fecal homogenate via oral gavage. Two weeks after the fecal community inoculation, mice were challenged with *C. difficile*. *C. difficile* clinical isolate 431 came from Carlson *et al*. which had previously been isolated and characterized (34, 35) and has recently been further characterized (36). Spores concentration were determined both before and after challenge (82). 10^3^ *C. difficile* spores were given to each mouse via oral gavage.

### Sample collection

Fecal samples were collected on the day of *C. difficile* challenge and the following 10 days. Each day, a fecal sample was collected and a portion was weighed for plating (approximately 30 mg) and the remaining sample was frozen at −20°C. Anaerobically, the weighed fecal samples were serially diluted in PBS, plated on TCCFA plates, and incubated at 37°C for 24 hours. The plates were then counted for the number of colony forming units (CFU) (83).

### DNA sequencing

From the frozen fecal samples, total bacterial DNA was extracted using MOBIO PowerSoil-htp 96-well soil DNA isolation kit. We amplified the 16S rRNA gene V4 region and sequenced the resulting amplicons using an Illumina MiSeq as described previously (84).

### Sequence curation

Sequences were processed with mothur(v.1.44.3) as previously described (84, 85). In short, we used a 3% dissimilarity cutoff to group sequences into operational taxonomic units (OTUs). We used a naive Bayesian classifier with the

Ribosomal Database Project training set (version 18) to assign taxonomic classifications to each OTU (86). We sequenced a mock community of a known community composition and 16s rRNA gene sequences. We processed this mock community with our samples to calculate the error rate for our sequence curation, which was an error rate of 0.19%.

### Toxin cytotoxicity assay

To prepare the sample for the activity assay, fecal material was diluted 1:10 weight per volume using sterile PBS and then filter sterilized through a 0.22-*μ*m filter. Toxin activity was assessed using a Vero cell rounding-based cytotoxicity assay as described previously (29). The cytotoxicity titer was determined for each sample as the last dilution, which resulted in at least 80% cell rounding. Toxin titers are reported as the log10 of the reciprocal of the cytotoxicity titer.

### Histopathology evaluation

Mouse cecal tissue was placed in histopathology cassettes and fixed in 10% formalin, then stored in 70% ethanol. McClinchey Histology Labs, Inc. (Stockbridge, MI) embedded the samples in paraffin, sectioned, and created the hematoxylin and eosin-stained slides. The slides were scored using previously described criteria by a board-certified veterinary pathologist who was blinded to the experimental groups (29). Slides were scored as 0-4 for parameters of epithelial damage, tissue edema, and inflammation and a summary score of 0-12 was generated by summing the three individual parameter scores.

### Statistical analysis and modeling

To compare community structures, we calculated Yue and Clayton dissimilarity matrices (*θ*_YC_) in mothur (87). We rarefied samples to 2,107 sequences per sample to limit uneven sampling biases. We tested for differences in individual taxonomic groups that would explain the outcome differences with LEfSe (88) in mothur. Remaining statistical analysis and data visualization was performed in R (v4.0.5) with the tidyverse package (v1.3.1). We tested for significant differences in *β*-diversity (*θ*_YC_) using the Wilcoxon rank sum test. We used Spearman’s correlation to identify which OTUs that had a correlation between their relative abundance and the histopathologic summary score. *P* values were then corrected for multiple comparisons with a Benjamini and Hochberg adjustment for a type I error rate of 0.05 (89). We built random forest models using the mikropml package (90) with relative abundance summed by taxonomic ranks from day 0 samples using mtry values of 1 through 10, 15, 20, 25, 40, 50, 100. The split for training and testing varied by model to avoid overfitting the data. To determine the optimal split, we tested splits (50%, 60%, 70%, 80%, 90% data used for training) to find the greatest portion of data that could be used to train the model while still maintaining the same performance for the training model as the model with the held-out test data. The toxin and moribundity models were trained with 60% of the data. The histopathologic score model was trained with 80% of the data. Lastly, we did not compare murine communities to donor community or clinical data because germ-free mice colonized with non-murine fecal communities have been shown to more closely resemble the murine communities than the donor species community (91). Furthermore, it is not our intention to make any inferences regarding human associated bacteria and their relationship with human CDI outcome.

### Code availability

Scripts necessary to reproduce our analysis and this paper are available in an online repository (https://github.com/SchlossLab/Lesniak_Severity_XXXX_2022).

### Sequence data accession number

All 16S rRNA gene sequence data and associated metadata are available through the Sequence Read Archive via accession PRJNA787941.

## Acknowledgements

Thank you to Sarah Lucas and Sarah Tomkovich for critical discussion in the development and execution of this project. We also thank the University of Michigan Germ-free Mouse Core for assistance with our germfree mice, funded in part by U2CDK110768. This work was supported by several grants from the National Institutes for Health R01GM099514, U19AI090871, U01AI12455, and P30DK034933. Additionally, NAL was supported by the Molecular Mechanisms of Microbial Pathogenesis training grant (NIH T32 AI007528). The funding agencies had no role in study design, data collection and analysis, decision to publish, or preparation of the manuscript.

---

**Figure S1.**
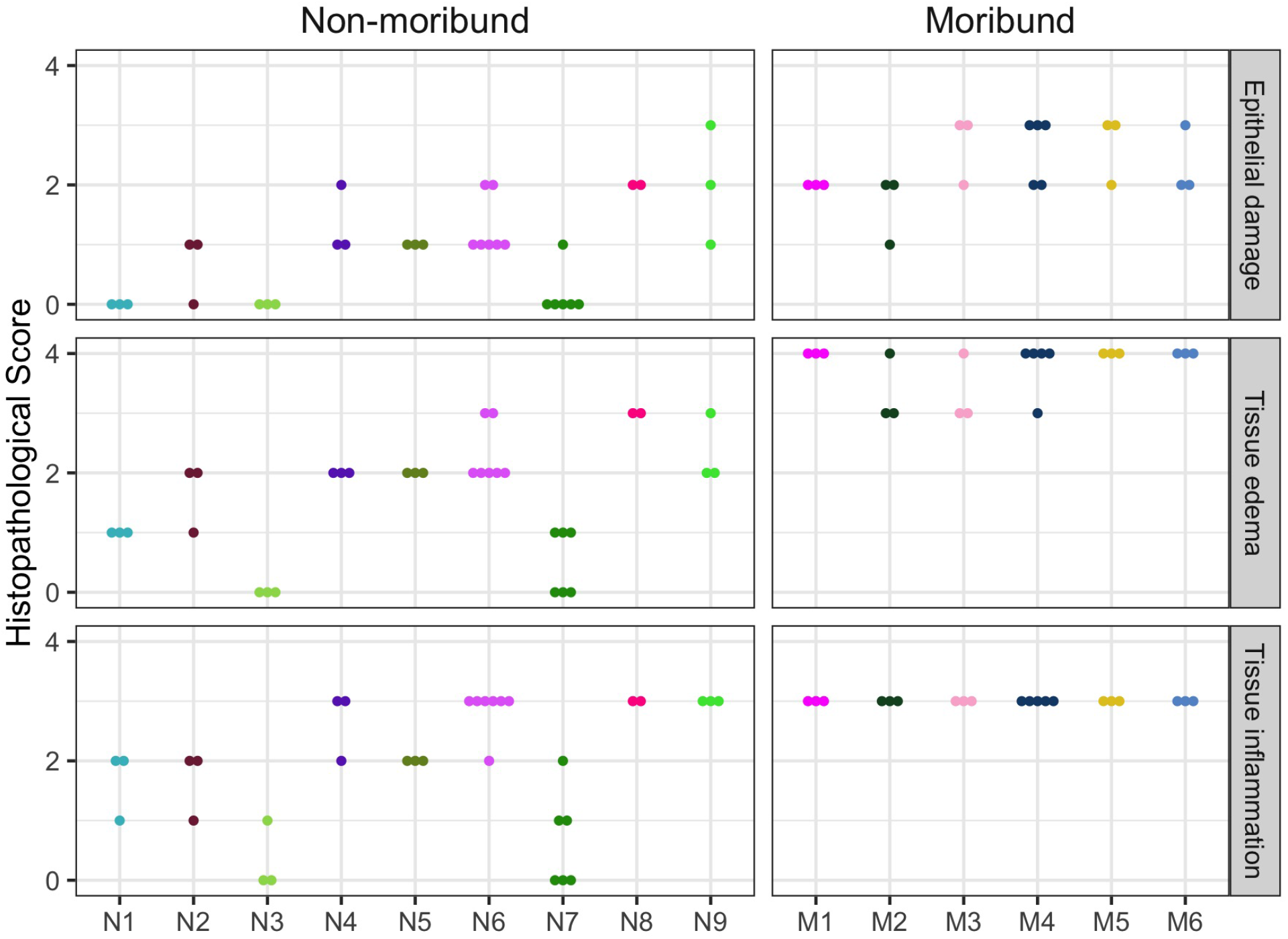
Histopathologic score of tissue damage at the endpoint of the infection. Tissue collected at the endpoint, either day 10 post-challenge (Non-moribund) or day mice succumbed to infection (Moribund), were scored from histopathologic damage. Each point represents an individual mouse. Mice (points) are grouped and colored by their human fecal community donor. Missing points are from mice that had insufficient sample for histopathologic scoring.

**Figure S2.**
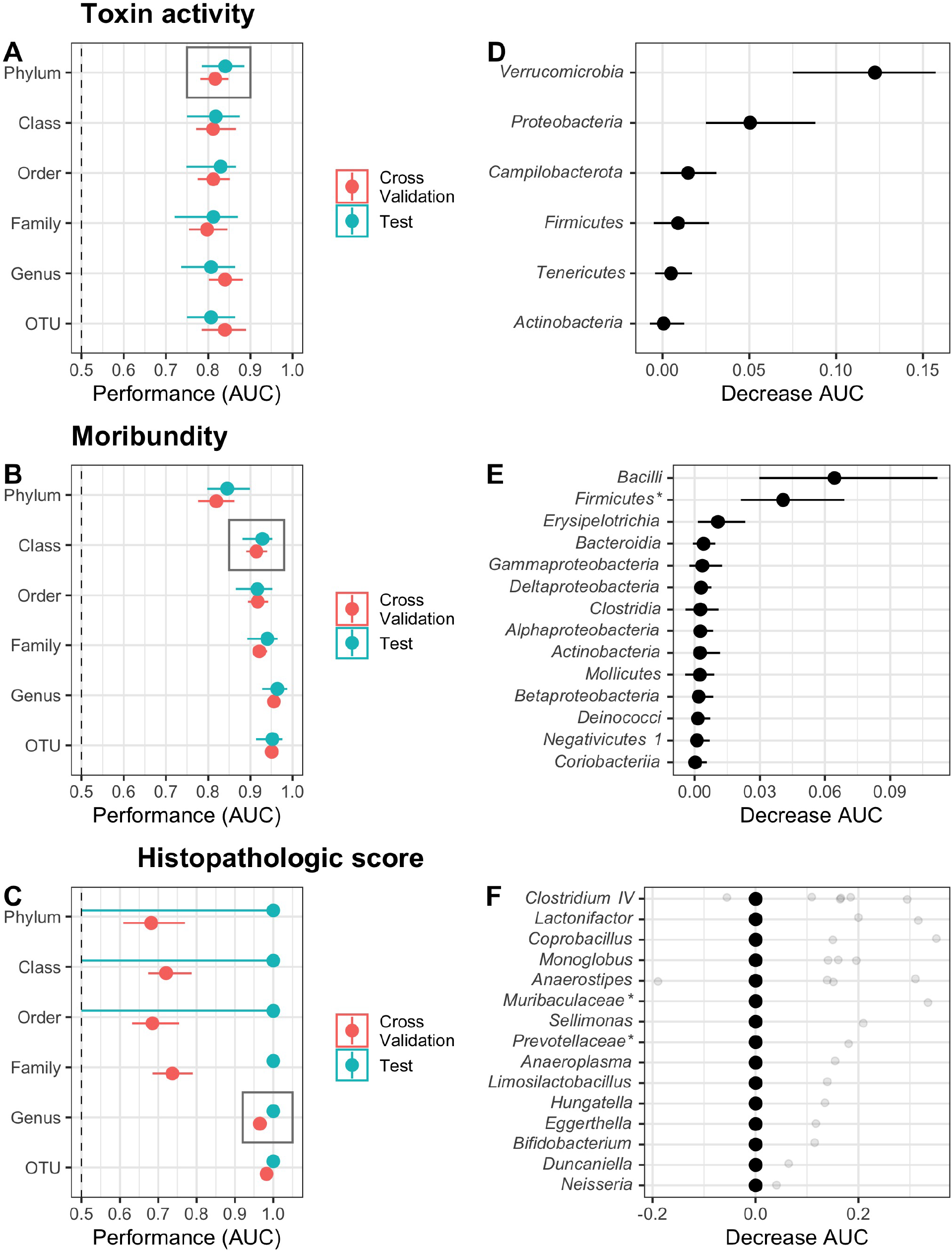
Random forest models predicted outcomes of the *C. difficile* challenge. (A-C) Taxonomic classification rank model performance. Relative abundance at the time of *C. difficile* challenge (Day 0) of the bacterial community members grouped by different classification rank were modeled with random forest to predict the infection outcome. The models used the highest taxonomic classification rank performed as well as the lower ranks. Black rectangle highlights classification rank used to model each outcome. (D-F) Model feature importance. Bacterial groups are ordered by their decrease in area under receiver-operator curve (AUC) when its relative abundances was permuted. Individual relative abundances were added to F since differences in AUC were outside the interquartile range. * indicates bacterial group was unclassified at lower taxonomic classification ranks. For all plots, median (solid points) and interquartile range (lines) are plotted. (A) Toxin production modeled which mice would have toxin detected during the experiment. (A) Moribundity modeled which mice would succumb to the infection prior to day 10 post-challenge. (C) Histopathologic score modeled which mice would have a high (score greater than the median score of 5) or low (score less than the median score of 5) histopathologic summary score. (D) Bacterial phyla which affected the performance of predicting detectable toxin activity when permuted. (E) Bacterial classes which affected the performance of predicting moribundity when permuted. (D) Bacterial genera which affected the performance of predicting histopathologic score when permuted.

